# Histone methyltransferase SUV39H1 regulates the Golgi complex via the nuclear envelope-spanning LINC complex

**DOI:** 10.1101/2023.03.13.532406

**Authors:** Miyu Nishino, Hiromasa Imaizumi, Yuhki Yokoyama, Jun Katahira, Hiroshi Kimura, Nariaki Matsuura, Miki Matsumura

## Abstract

Cell motility is related to the higher-order structure of chromatin. Stimuli that induce cell migration change chromatin organization; such stimuli include elevated histone H3 lysine 9 trimethylation (H3K9me3). We previously showed that depletion of histone H3 lysine 9 methyltransferase, SUV39H1, suppresses directional cell migration. However, the molecular mechanism underlying this association between chromatin and cell migration remains elusive. The Golgi apparatus is a cell organelle essential for cell motility. In this study, we show that loss of H3K9 methyltransferase SUV39H1 but not SETDB1 or SETDB2 causes dispersion of the Golgi apparatus throughout the cytoplasm. The Golgi dispersion triggered by SUV39H1 depletion is independent of transcription, centrosomes, and microtubule organization, but is suppressed by depletion of any of the following three proteins: LINC complex components SUN2, nesprin-2, or microtubule plus-end-directed kinesin-like protein KIF20A. In addition, SUN2 is closely localized to H3K9me3, and SUV39H1 affects the mobility of SUN2 in the nuclear envelope. Further, inhibition of cell motility caused by SUV39H1 depletion is restored by suppression of SUN2, nesprin-2, or KIF20A. In summary, these results show the functional association between chromatin organization and cell motility via the Golgi organization regulated by the LINC complex.

## Introduction

Nuclear deformation occurs during cell migration [1], and several lines of evidence including our data show a link between chromatin structure and cell migration. The dynamic reorganization of the chromatin fiber is an early event in the cellular response to migration cues [2-4]. In particular, cell migration cues induce elevated histone H3 lysine 9 trimethylation (H3K9me3), histone H4 lysine 20 monomethylation (H4K20me1), and histone H3 lysine 27 trimethylation (H3K27me3) [5]. Activated cell migration occurs concomitantly with up-regulated H3K9me3 [6], and H3K9-specific histone methyltransferase, SUV39H1, is involved in regulating cell migration [7]. SUV39H1 specifically di- and tri-methylates H3K9 and modulates chromatin dynamics, telomere length, heterochromatin organization, chromosome segregation, and mitotic progression [8-10]. However, the molecular link between chromatin organization and cell migration is still unclear.

The mammalian Golgi apparatus is the central organelle within the secretory pathway and the formation of cell polarity. In most vertebrate cells, individual Golgi stacks are laterally connected in a juxtanuclear array, termed the Golgi ribbon, in close association with the centrosome, which is localized in the juxta nuclear area and functions as the MT-organizing center. During cell migration, physical contact between the Golgi and the centrosome allows for a coordinated re-localization of both structures [11]. Centrosome- and Golgi-derived MTs cooperate with actin filaments to maintain the Golgi structure [12]. The structure of the Golgi complex is highly dynamic, and the Golgi morphology is influenced by cellular contexts such as cell migration or differentiation, which correlates with nuclear functions. However, mechanistic associations between the architecture of the mammalian Golgi complex and nuclear function is still puzzling.

The linker of nucleoskeleton and cytoskeleton (LINC) complex, a multifunctional nuclear envelope protein complex, consists of the inner nuclear membrane-spanning Sad1 and UNC-84 (SUN) domain-containing proteins, SUNs, and the outer nuclear membrane-spanning Klarsicht, ANC-1, and Syne homology (KASH) domain-containing proteins, nesprins. SUNs and nesprins associate with each other in the lumen of nuclear envelope [13]. SUN proteins interact with nuclear components such as lamins and chromatin, whereas nesprins associate with cytoskeletal components including microtubule (MT) motors and actin filaments [14]. Thus, the LINC complex directly connects the cytoskeleton and nucleoskeleton to form a scaffold for diverse functions including nuclear migration, shaping, and positioning [15], maintenance of centrosome positioning and its nuclear connection [16, 17], mechanotransduction [18, 19], DNA repair [20, 21], nuclear membrane spacing [22], cell migration [23, 24], moving chromosomes within the nucleus during meiosis [25, 26], and Golgi organization [27]. In addition, nesprin and SUN proteins are essential for neurological and muscular development in mice [16, 28], and mutations in nesprin- and SUN-encoding genes in humans have been shown to cause or be linked to muscular dystrophies, ataxia, progeria, and multiple cancers [29, 30].

Mammalian somatic cells mainly express two SUN genes, SUN1 and SUN2. We recently showed that depletion of SUN1 leads to Golgi dispersion with maintenance of ministacks [27]. This dispersion of the Golgi complex requires the function of plus end-directed MT motor KIF20A in conjunction with the SUN2/nesprin-2 containing LINC complex. In addition, SUN2 associates with KIF20A, possibly via nesprin-2. In this study, we investigated the association between chromatin organization and cell migration and found that the nuclear status such as the histone modification links to the Golgi organization through the LINC complex and affects cell migration.

## Results

### SUV39H1 is required to maintain the Golgi complex architecture

Cell migration cues change the global chromatin organization including elevated H3K9me3 [31], and depletion of SUV39H1 affects directional cell migration [7]. However, the molecular link between chromatin and migratory activity has not been determined. Because proper Golgi organization is required for directional cell migration, the effects of a histone lysine methyltransferase inhibitor, chaetocin, on the Golgi architecture was examined. While the Golgi complex is generally positioned between the nucleus and the leading edge during directional migration (Fig 1A, left), the drug induced a scattering of the Golgi complex throughout the cytoplasm (Fig 1A, right). The Golgi complex was detected using an antibody specific for the cis-Golgi marker GM130 (Golgi matrix protein 130). Quantified analysis showed a randomly oriented Golgi complex in the chaetocin-treated cells (Fig 1B). The dispersal of the Golgi complex triggered by the histone methyltransferase inhibitor was not specific to the MDA-MB-231 cells but was also observed in other cells such as HeLa (S2D, Fig). HeLa cells express at least three H3K9-specific histone methyltransferases including SETDB1, SETDB2, and SUV39H1. To assess the specificity of the drug, these H3K9me3 methyltransferases were depleted using siRNA. Depletion of either SETDB1 or SETDB2 did not affect the morphology of the Golgi complex, while the expression level of SETDB1 and SETDB2 mRNA were reduced by 90% as determined by real-time PCR (RT-PCR, S1A and S1B Fig). In contrast, the SUV39H1-depleted cells showed drastically dispersed Golgi complexes throughout the cytoplasm (Fig 1C and S1A Fig for a low magnification field of view). The reduction of SUV39H1 mRNA was confirmed by RT-PCR (Fig 1D). In this study, a pool of three siRNAs against SUV39H1 was used because similarly scattered Golgi complexes were observed in each of the three siRNA-transfected cells (S1C Fig). Upon depletion of SUV39H1, the GM130 positive area increased more than twofold (Fig 1E), and the impaired Golgi structure was rescued by an expression of siRNA-resistant mouse SUV39H1 (Fig 1F). In addition, the Y67A SUV39H1 mutant, which contains a point mutation in the chromodomain and impairs enzyme activity [32, 33], also induced the Golgi dispersal. This dominant negative effect indicates that proper activity of the SUV39H1 is required for the morphological integrity of the Golgi complex (Fig 1G). Therefore, these results demonstrate that SUV39H1 specifically associates with architecture of the Golgi complex.

**Fig 1.**
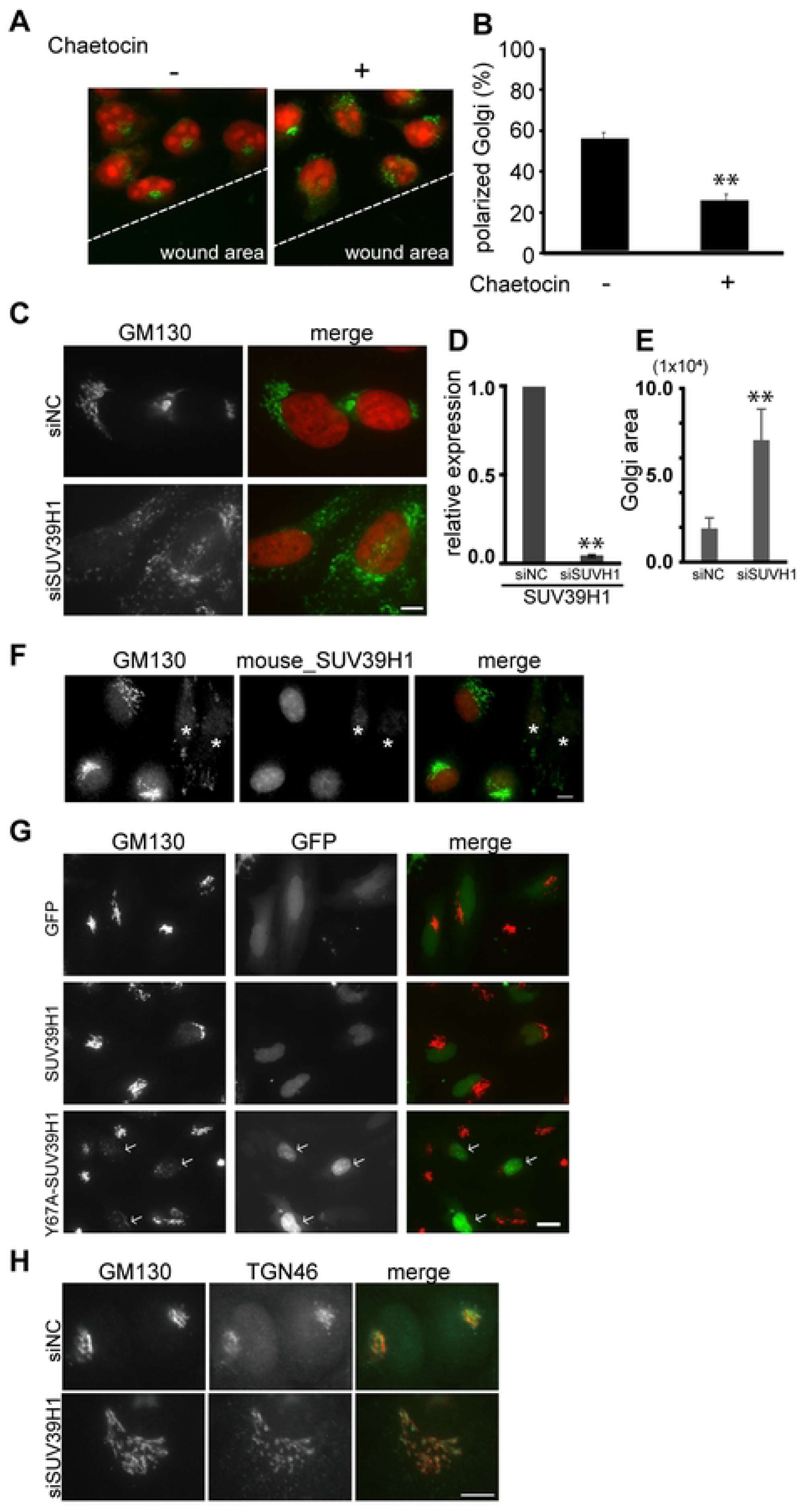
SUV39H1 associates with the architecture of the Golgi complex. **A.** Confluent breast cancer cells, MDA-MB-231, which show active cell migration, were scratched with or without a histone methyltransferase inhibitor, chaetocin. After 5 h incubation, the cells were fixed and stained with anti-GM130 mAb, a cis-Golgi marker (green). The nucleus was counterstained with propidium iodide, PI (red). **B.** The cells with the Golgi complex located within the front quarter of the cell, facing the wound area, were considered “polarized”; the cells with the Golgi complex located in the rear three-quarters of the cell body, away from the wound, were considered “nonpolarized”. The ratio (%) ±SE of the cells containing the polarized Golgi complex is indicated (n > 100). ** P < 0.01, compared with the chaetocin-untreated cells. **C.** HeLa cells were transfected with siRNA against SUV39H1 (siSUV39H1) or negative control siRNA (siNC). After 48 h incubation, the cells were stained with DAPI (red) and anti-GM130 mAb (green). Bar, 10 μm. **D.** mRNA was obtained from the SUV39H1-depleted cells and analyzed by RT-PCR. ** P < 0.01, compared with the control cells. **E.** Quantification of GM130 labeled Golgi complex area per cell. ** P < 0.01, compared with control cells. **F.** Cells were transfected with siSUV39H1. Then, GFP-tagged mouse SUV39H1 was transfected. The cells were fixed and GFP and GM130 were detected. Asterisks show mouse_SUV39H1 untransfected cells. Bar, 10 μm. **G.** Cells were transfected with GFP, GFP-tagged SUV39H1 or GFP-tagged Y67A-SUV39H1. GFP and GM130 were then detected. Arrows show GFP-tagged Y67A-SUV39H1 transfected cells. **H.** Cells were transfected with siSUV39H1 or siNC. The cells were then stained with anti-GM130 pAb and anti-TGN46, a trans-Golgi marker protein, mAb. Bar, 10 μm.

The dispersed Golgi complex was observed in greater than 90% of the SUV39H1-depleted cells (S1A Fig), implying that this phenomenon is not cell cycle dependent. In addition, the dispersed GM130 molecules in the SUV39H1-depleted cells were situated in close proximity to other Golgi complex marker proteins, TGN46 (a trans-Golgi marker protein) and Golgi 58K protein (a medial-Golgi marker protein) (Fig 1H and S1D Fig), suggesting that these dispersed Golgi complexes maintain mini-stacks that remain competent for posttranslational processing and protein secretion [34]. Indeed, a newly synthesized plasma membrane protein, amphiregulin, was transported to the plasma membrane in the SUV39H1-depleted cells (S2A Fig).

A collapsed Golgi complex has been observed in a variety of conditions such as prevention of MT dynamics or inhibition of ER-to-Golgi protein delivery. However, SUV39H1 depletion affects neither the MT organization (S2B Fig) nor ER-to-Golgi transport (S2A Fig), indicating that either disruption of MT organization or suppression of ER-to-Golgi transport is not a reason for the dispersed Golgi complex in the SUV39H1-depleted cells. Moreover, the centrosome in the SUV39H1-depleted cells remains in the perinuclear region, similarly to the control cells (S2C Fig). This result implies that Golgi dispersion triggered by SUV39H1 depletion is independent of the centrosome.

Furthermore, 5,6-dichloro-1-β-ribofuranosylbenzimidazole (DRB), a transcription inhibitor, has no effect on Golgi architecture, and the disruption of Golgi architecture by chaetocin treatment occurs even in the presence of DRB (S2D Fig). H3K9 methylation is a constitutive heterochromatin mark, and induction of migration leads to a higher increase in H3K9me3 signals at repetitive elements than at protein-coding genes [5]. These lines of evidence indicate that the Golgi complex scattering in the SUV39H1-depleted cells is independent of gene expression. We therefore decided to examine the possibility that chromatin organization directly links to Golgi complex integrity.

### KIF20A is required for the Golgi dispersal observed in the SUV39H1-depleted cells

Juxtanuclear positioning of the Golgi complex is the balanced outcome of minus- and plus-end-directed MT-dependent motors. A plus-end-directed kinesin family member, KIF20A (also known as mitotic kinesin-like protein 2, MKlp2 or Rab6-interacting kinesin-like protein, *RAB6KIFL*), is localized to the Golgi complex in the interphase cells and plays a role in the dynamics of the Golgi complex [35, 36, 37]. In cells expressing large amount of KIF20A, the Golgi are scattered into small structures throughout the cytoplasm [35]. Thus, to assess a commitment of KIF20A to the Golgi dispersal caused by SUV39H1 depletion, a KIF20A inhibitor, paprotrain, was used. The drug suppressed the Golgi dispersal in the SUV39H1-depleted cells (Fig 2A, right), whereas the drug did not affect Golgi morphology in the control cells (Fig 2A, left). Next, to confirm the function of KIF20A in the Golgi dispersion, KIF20A was depleted together with SUV39H1 using siRNA. This treatment also markedly suppressed the Golgi dispersal caused by SUV39H1 depletion (Fig 2B). The area of the Golgi complex in SUV39H1 and KIF20A double knock down cells was reduced to a similar level as the control cells (Fig 2C). These results indicate that KIF20A functions in Golgi scattering in the SUV39H1-depleted cells. Note that because KIF20A functions in both Golgi maintenance and mitosis exit, its depletion inhibited cytokinesis and results in the accumulation of multinucleated cells [38] (Fig 2B).

**Fig 2.**
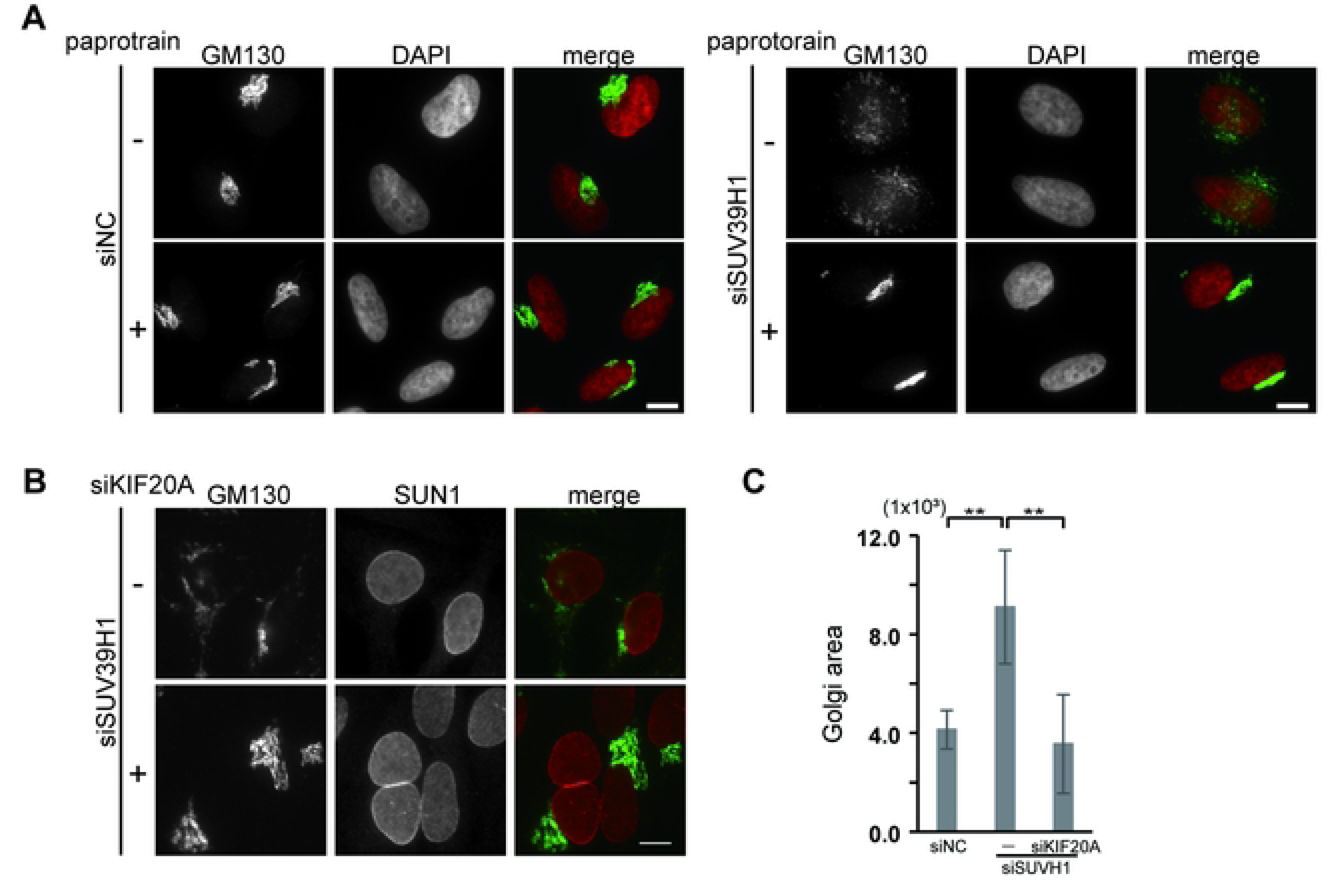
KIF20A is required for the Golgi dispersal observed in the SUV39H1-depleted cells. **A.** Cells were transfected with siSUV39H1 and incubated for 36 h. Paprotrain was then added. After 12 h, the cells were fixed and stained with anti-GM130 mAb. Bar, 10 μm. **B.** Cells were transfected with siRNA against KIF20A (siKIF20A) and/or siSUV39H1. After 48 h incubation, the cells were fixed and stained with anti-GM130 mAb and anti-SUN1 pAbs to show the nuclei. **C.** Quantification of GM130 labeled Golgi complex area per cell in the SUV39H1- and/or KIF20A-depleted cells. ** P < 0.01, compared with control cells.

### SUN2/nesprin-2 LINC complex functions in the Golgi dispersal caused by SUV39H1 depletion

To investigate how the nuclear event, i.e., SUV39H1 depletion, influences cytoplasmic organelle integrity, we focused on the LINC complex, which is composed of integral inner and outer nuclear membrane proteins, SUN and nesprin, respectively, because we recently showed that the SUN2/nesprin-2 LINC complex plays a role in Golgi dispersal together with KIF20A [27]. We first verified the LINC complex requirement in the Golgi scattering in SUV39H1-depleted cells using a dominant negative KASH construct (DN-KASH), which broadly interferes with nesprin–SUN interaction [39]. Expression of DN-KASH suppressed Golgi dispersal in the SUV39H1-depleted cells compared with untransfected cells in the same field (Fig 3A, arrows indicate transfected cells). The quantified data indicate the requirement of the LINC complex for Golgi dispersal induced by SUV39H1 depletion (Fig 3B).

**Fig 3.**
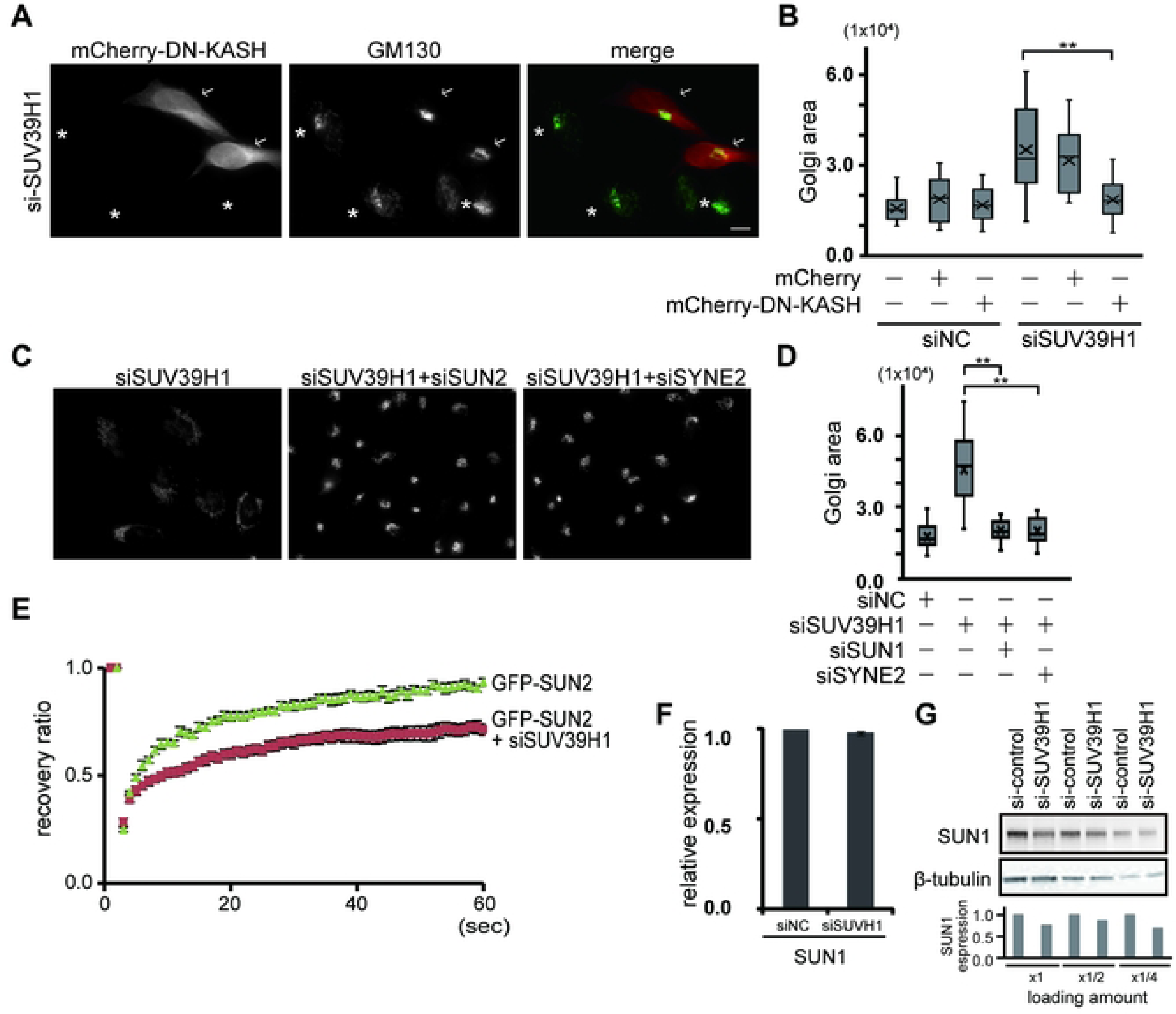
SUN2/nesprin-2 LINC complex functions in the Golgi dispersal caused by SUV39H1 depletion. **A.** Cells were transfected with siSUV39H1. After 24 h incubation, the cells were again transfected with mCherry tagged DN-KASH. The cells were then fixed and stained with anti-GM130 mAb (green). Arrows and asterisks show mCherry tagged DN-KASH transfected and untransfected cells, respectively. **B.** Box-and-whiskers plots represent the GM130-labeled Golgi complex area. ** P < 0.01. **C.** Cells were transfected with siSUV39H1 and siSYN2 or siSUN2, which targets nesprin-2 and SUN2, respectively. The cells were then stained with anti-GM130 mAb. Bar, 10 μm. **D.** Box-and-whiskers plots represent the GM130-labeled Golgi complex area in the indicated cells. ** P < 0.01. **E.** To investigate SUN1 mobility, cells were transfected with siSUV39H1 or siNC; on the following day, the cells were transfected with GFP-tagged SUN2. FRAP analysis was then performed (n ≥ 10). Note that there are no significant differences in SUN2 mobility between siNC-transfected and -untransfected cells. The results are presented as means ± SEM. **F.** Cells were transfected with siSUV39H1, and SUN1 mRNA expression was analyzed by RT-PCR. **G.** Cells were transfected with siSUV39H1 and SUN1, and β-tubulin protein expression was analyzed by western blotting. SUN expression was represented as the amount of SUN1 in siNC-transfected cells as 1.

Next, to more directly analyze the involvement of SUN2 or nesprin-2, these proteins were depleted together with SUV39H1 depletion. Depletion of either SUN2 or nesprin-2 suppressed the Golgi scattering upon the loss of SUV39H1. The Golgi were localized in the perinuclear region with a normal morphology in cells lacking SUN2 or nesprin-2 along with SUV39H1 (Fig 3C). The quantified data suggest that the LINC complex composed of SUN2 and nesprin-2 plays a role in the dispersal of the Golgi complex caused by SUV39H1 depletion (Fig 3D).

Then, to investigate the possibility that SUN2 is incorporated into the LINC complex in the SUV39H1-depleted cells, we validated the mobility of SUN2 at the nuclear envelope in the presence and absence of SUV39H1 through fluorescence recovery after photobleaching (FRAP) analysis. Unlike previously shown mouse SUN2 data [40], human SUN2 is more mobile, and SUV39H1 depletion reduced SUN2 mobility, implying the formation of a SUN2/nesprin-2 LINC complex in the SUV39H1-depleted cells (Fig 3E).

Because our previous results show that the SUN2/nesprin-2 LINC complex functions in Golgi dispersion when SUN1 is absent [27], alterations in the Golgi architecture in SUV39H1-depleted cells might be mediated by loss of SUN1 expression. However, contrary to our expectation, SUN1 mRNA expression was not influenced by SUV39H1 depletion (Fig 3F), and protein expression is detectable but reduced around 80% (Fig 3G and S3 Fig). Therefore, these results indicate two things: first, SUN2/nesprin-2 and KIF20A contribute to the Golgi dispersal in the SUV39H1-depleted cells, and second, the Golgi dispersion in the SUV39H1-depleted cells is not due to SUN1 deficiency.

### H3K9me3 and SUN2 localize in close proximity

SUV39H1 directs trimethylation of H3K9, which is frequently found in heterochromatic regions including the nuclear periphery. To examine the positional relationship between H3K9me3 and SUN proteins, we performed a proximal ligation (PLA) assay and found a close localization of H3K9me3 with SUN2 (Fig 4A). In addition, SUV39H1 depletion decreased the signal numbers obtained by the association between SUN2 and H3K9me3 (Fig 4B). The reduction was confirmed by quantification (Fig 4C). These results indicate that SUN2 and H3K9me3 are localized in close proximity.

**Fig 4.**
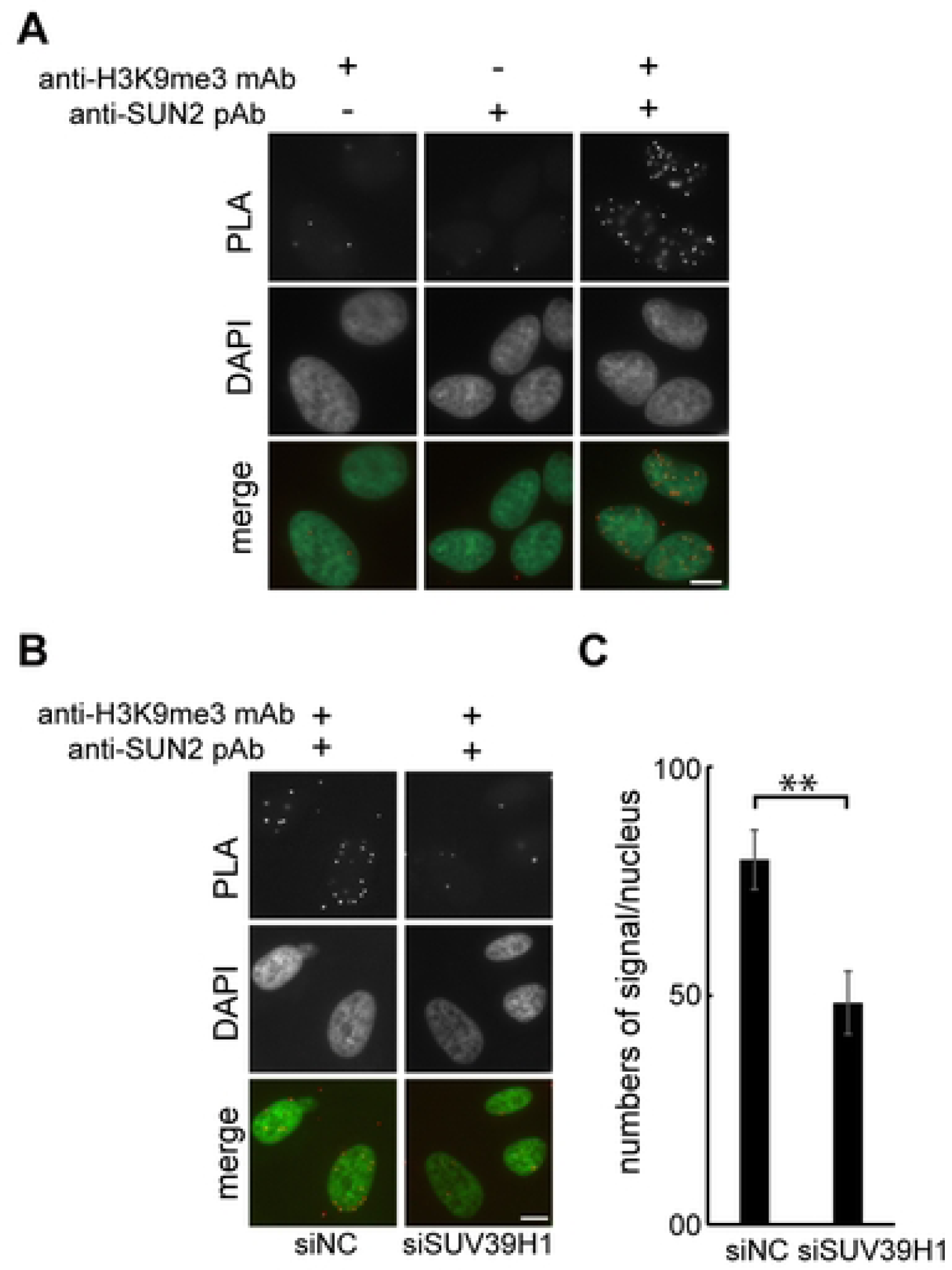
H3K9me3 and SUN2 localize in close proximity. **A.** Cells were fixed and incubated with the indicated antibodies, and signals were detected with PLA and DAPI. **B.** Cells were transfected with siSUV39H1 or siNC. Fixed cells were then incubated with anti-H3K9me3 mAb and anti-SUN2 pAb, and signals were detected with PLA and DAPI. **C.** The numbers of PLA signals between H3K9me3 and SUN2 were counted in the cells transfected with siSUV39H1 or in the siNC-transfected cells. The results are presented as the mean number in a nucleus ± SE. ** P < 0.01.

### Loss of SUN2, nesprin-2, or KIF20A restored suppressed cell motility in the SUV39H1-depleted cells

The Golgi complex is an essential organelle for cell motility. SUV39H1 depletion induces dispersion of the Golgi complex (Fig 1) and suppresses cell motility [7]. In addition, the Golgi dispersion induced by the loss of SUV39H1 is suppressed by depletion of either SUN2, nesprin-2 or KIF20A. Based on these results, to investigate whether the impaired Golgi complex is the reason for the suppressed cell motility in the SUV39H1-depleted cells, SUN2, nesprin-2 or KIF20A was knocked down in conjunction with SUV39H1 depletion and a wound healing assay was performed.

Wounds generated by scratching gradually closed over 18 h in the control cells, and SUV39H1 depletion impaired wound closure over the same time period as previously reported [7]. In contrast, double depletion of either SUN2, nesprin-2, or KIF20A but not SUN1 along with SUV39H1 protein rescued the suppressed wound closure (Fig 5A). The quantified data support this conclusion (Fig 5B). These results indicate that chromatin organization associates with cell motility through the Golgi organization, which is regulated by the LINC complex.

**Fig 5.**
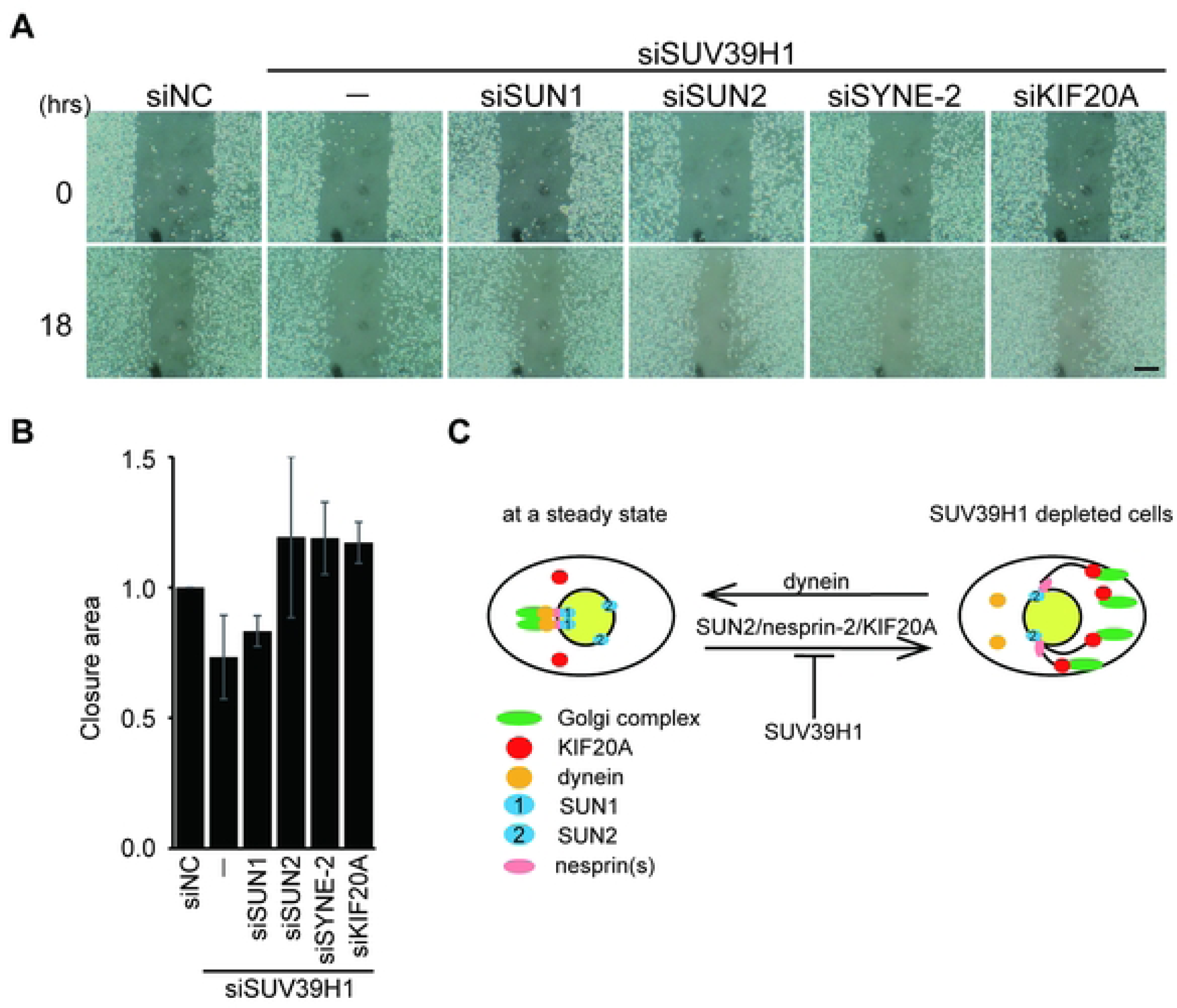
The dispersed Golgi complex is responsible for the suppression of cell motility resulting from loss of SUV39H1. **A.** MDA-MB-231 were transfected with the indicated siRNA. Confluent cells were scratched to create a model wound. Phase-contrast images 18 h after scratching to create the wound. **B.** Cell motility was quantified using phase-contrast images in A. Eighteen hours after wounding, the area of unclosed wounds was measured. The wound closures are shown relative to the area of unclosed wounds in siNC-transfected cells (means ± SE). Bar, 300 μm. ** P < 0.01, compared with siSUV39H1 transfected cells. **C.** A model in which SUV39H1 plays a critical role in maintaining the Golgi apparatus through the function of the LINC complex and KIF20A. In a steady state, i.e., in the presence of SUV39H1, SUV39H1 negatively regulates SUN2/nesprin-2 LINC complex formation (left), and the cells show the perinuclear localization of the Golgi complex. Upon the depletion of SUV39H1 function, the suppression of SUN2/nesprin-2 LINC complex formation is restored, and thus SUN2/nesprin-2/KIF20A function is activated, resulting in the scattered Golgi complex (right).

## Discussion

The structure and positioning of the Golgi complex dramatically changes during several cellular processes such as cell migration and differentiation. These processes correlate with nuclear functions, while the molecular mechanism underlying the correlation between Golgi morphology and nuclear functions has not been determined. In this study, we showed that histone H3 lysine 9 methyltransferase, SUV39H1, is required for a proper Golgi structure. Depletion of SUV39H1 leads to the scattering of the Golgi complex, presumably through the chromatin structure rather than through affecting gene expression. In addition, the LINC complex components SUN2, nesprin-2, and a plus-end-directed MT motor protein, KIF20A, is required for the Golgi fragmentation caused by loss of SUV39H1. Based on these findings, we propose a model in which SUV39H1 plays a critical role in maintaining the Golgi apparatus through the function of the LINC complex and KIF20A (Fig 5C). SUN2 physically interacts with KIF20A [27]. Loss of KIF20A, SUN2, or nesprin-2 is sufficient to suppress Golgi fragmentation caused by SUV39H1 depletion (Fig 2 and 3). Thus, the SUN2/nesprin-2 LINC complex and KIF20A cooperatively function in Golgi dispersion. In a steady state, the Golgi complex is localized at the nuclear periphery with a dynamic equilibration of minus- and plus-end-directed MT-dependent motor activity. The perinuclear positioning of the Golgi complex is mainly maintained through the action of MT minus-end-directed motor dynein. In contrast to the dynein function, KIF20A moves toward the plus-ends of MTs and exerts force on the Golgi complex. In the presence of SUV39H1, KIF20A and/or SUN2/nesprin-2 activity is suppressed, and thus the Golgi complex is localized at the nuclear periphery. PLA analysis shows a proximate localization of H3K9me3 and SUN2 (Fig 4). The results of FRAP analysis imply that incorporation of SUN2 into the LINC complex is suppressed by SUV39H1 (Fig 3). These findings indicate that H3K9me3 mediated by SUV39H1 negatively regulates SUN2/nesprin-2 LINC formation, and the Golgi complex accumulates in the juxtanuclear area (Fig 5C, left). In contrast, upon SUV39H1 depletion, the SUN2/nesprin-2 LINC complex forms and KIF20A is activated, resulting in Golgi dispersion throughout the cytoplasm (Fig 5C, right). The following three lines of evidence support that LINC complex formation is biased toward SUN1 inclusion and SUN2/nesprin-2 LINC complex formation is likely to be suppressed in a steady state: (i) the amount of SUN1 and SUN2 is much higher than nesprins-2; (ii) SUN2 binding to KASH is weaker than SUN1; and (iii) SUN1 is more efficiently incorporated into LINC complexes than SUN2 under normal growth conditions [40, 41]. In addition to the above evidence, we show that H3K9me3 is localized in the vicinity of SUN2 and inhibits its incorporation into the LINC complex, supporting the biased incorporation. In conclusion, we show a potential role of chromatin organization in regulating Golgi integrity through the LINC complex.

Heterochromatin reorganization upon activation of cell migration has been observed in a variety of cells including T cells, bone marrow-derived mesenchymal stem cells, and cancer cells such as melanoma [1-7, 42, 43]. For instance, CD4^+^ T cells, whose motility is activated by vascular cell adhesion molecule 1 (VCAM1), increase the H3K9me2/3 level and the resistance of the genome to cleavage by DNase I and MNase upon induction of migration [42]. Migration cues elevate H3K9me3, H3K27me3, and H4K20me1 in melanoma cells [2, 3, 5]. In colon cancer tissue, invasive but not superficial regions show an increased H3K9me3 level [7]. In addition, heterochromatin affects the cell migration rate. Depletion of either a catalytic subunit of the H3K27 methyltransferase complex PRC2, EZH2, or H3K9 methyltransferases including G9a, SUV39H1, and SETDB1/2 reduces the migration rate of various cell types [reviewed in 31]. Heterochromatin supports the migration process by contributing both the mechanical property of the nucleus as well as transcriptional regulation within the nucleus [31]. Increased heterochromatin enhances nuclear rigidity in a manner that allows faster cell migration in 3D environments. Besides these two functional significances of chromatin in cell migration (mechanical and transcriptional functions), our results in this study demonstrate another role of chromatin: its posttranslational modification regulates Golgi complex integrity via the LINC complex. This molecular link between SUV39H1 and the Golgi complex can explain previous observations, the correlations between cell migratory activities and chromatin structure [1-3, 7].

Disruption of the Golgi complex induces neurodegenerative diseases, e.g., loss of GM130 causes Golgi disruption in cerebellar Purkinje neurons and induces ataxia in mice [44]. Conversely, disruption of the Golgi architecture and function are widely observed in neurodegenerative diseases including Alzheimer’s disease and Parkinson’s disease [45, 46], and various cancer tissues [47, 48]. Global changes in the epigenetic landscape are a hallmark of cancer. In addition, the LINC complex has been shown to function in neurogenesis [16, 49]. Sun1 is selectively expressed in Purkinje cells in the cerebellum and Sun1-knockout mice show a marked decrease in cerebellar size, and reduced foliation of the cerebellar cortex, which causes cerebellar ataxia [49]. The Golgi complex is disorganized in Purkinje cells in SUN1-knockout mice [27]. In this study, we showed that SUV39H1 depletion induces fragmentation of the Golgi complex via the LINC complex. Therefore, it is possible that the impaired Golgi complex induced by altered chromatin organization associates with neurodegenerative diseases and cancer.

In summary, we showed that SUV39H1 plays a role in maintaining Golgi complex integrity through SUN2/nesprin-2 LINC complex formation. In addition, our data show that the LINC complex relays nuclear status such as histone modification to the Golgi complex. This is in the opposite direction to previously known functions of the LINC complex, such as mechanotransduction [18, 19] or meiotic telomere clustering [25, 26] in which the LINC complex transfers cytoplasmic forces or signaling from the cytoskeleton into the nucleus. Therefore, the LINC complex bi-directionally communicates and coordinates between the cytoplasm and nucleus.

## Materials and Methods

### Antibodies and solutions

Mouse anti-GM130 mAb (clone 35) was from Becton Dickinson and Company, BD (Franklin Lakes, NJ, USA) and used at a dilution of 1:200. Because GM130 is also known as GOLGA2, anti-GOLGA2 pAb (HPA021799) was purchased from Sigma Aldrich (St. Louis, MO, USA) and used at a dilution of 1:200. Mouse anti-Golgi 58K protein (formimidoyltransferase-cyclodeaminase) mAb (G2404) was from Sigma-Aldrich and used at a dilution of 1:50∼100. Rabbit anti-SUN1 pAb (HPA008346) and mouse anti-β-tubulin mAb (T4026, clone TUB2.1) were also from Sigma-Aldrich. Both of them were used at a dilution of 1:200 for immunofluorescent microscopy and at a dilution of 1:1000∼2000 for western blotting. Sheep anti-TGN46 pAb (AHP500G. 1:50) was from Serotec (Oxford, UK). Anti-SUN2 pAb (#06-1038) was purchased from Millipore (Rahway, NJ, USA) and used at a dilution of 1:200. Rabbit anti-KIF20A pAb (ab70791) was obtained from Abcam (Cambridge, UK). Rat anti-HA mAb (3F10) and rabbit anti-GFP pAb (code No. 598) were from Roche (Basel, Switzerland) and Medical and Biological Laboratories (Aichi, Japan), respectively. Mouse anti-H3K9me3 mAb was previously described [7, 50]. Mouse anti-centrin mAb (clone H20-5) was purchased from Millipore and used at a dilution of 1:50∼100. Chaetocin [51] and DRB were from Sigma-Aldrich. Paprotrain, KIF20A inhibitor, was purchased from Sigma Aldrich and used at a final concentration of 20 μM [52].

### Cell culture, transfection, and immunofluorescence

HeLa cells were obtained from the JCRB cell bank (Japanese Collection of Research Bioresources, Japan) and grown in Dulbecco’s modified Eagle’s medium, low glucose (Fujifilm Wako Chemical, Osaka, Japan) supplemented with 10% fetal calf serum at 37°C in a 10% CO_2_ atmosphere. The human metastatic human breast cancer cell MDA-MB-231 (ATCC HTB-26) was cultured as previously described [53]. Transfection was performed using Lipofectamine 2000 (Thermo Fisher Scientific, Waltham, MA, USA) or GeneJuice (Merck Millipore, Rahway, NJ, USA) in accordance with the manufacturers’ instructions. Unless stated otherwise, 17–20 hours after transfection, cells were fixed with 4% paraformaldehyde and immunofluorescence was performed as described previously [54] using appropriate primary and secondary antibodies (Jackson ImmunoResearch Laboratories, West Grove, PA, USA), and coverslips were mounted using Prolong Gold Antifade Reagent with 4′,6-diamidino-2-phenylindole (DAPI) (Life Technologies, Carlsbad, CA, USA). Cells were viewed using an Olympus IX81 or Olympus BX53 epifluorescence microscope. The proximal ligation assay kit (PLA) was purchased from Nacalai Tesque, Inc. (Kyoto, Japan) and performed according to the manufacturer’s protocol. To quantify the signals, images were captured and signals in a nucleus were manually counted (n > 25). Experiments were repeated at least three times.

### Short interfering RNA (siRNA)-mediated knockdown

Predesigned siRNA pools (siGenome SMARTpool; each siRNA pool contains four siRNAs) against SETDB1, SETDB2, SUN1 (UNC84A), SUN2 (UNC84B), KIF20A, and SYNE2 (nesprin-2) were obtained from Thermo Fisher Scientific, and siRNA oligos against SUN1 and SUN2 were obtained from Nippon Gene (Tokyo, Japan). siRNA oligos against SUV39H1 were obtained from Nippon Gene. The sequences of the siRNA pools targeting SUN1, SUN2, KIF20A, SUV39H1, and SYNE2 were described previously [7, 23, 27] and others are listed in Table S1A. Listed siNC, negative control siRNA (siGenome nontargeting siRNA pool, a mixture of four nontargeting siRNAs), was obtained from Thermo Fisher Scientific and Nippon Gene. Cells were transfected with siRNAs Lipofectamine RNAi MAX (Invitrogen, Waltham, CA, USA) in accordance with the manufacturer’s protocol. Cells were fixed or harvested 48 h after transfection unless otherwise stated. All siRNAs were used at a final concentration of 10 nM. To verify knockdown efficiency, the expression of the corresponding mRNA was detected by RT-PCR using the appropriate primer sets (Table S1B).

### Fluorescence recovery after photobleaching (FRAP) analysis

For the FRAP experiments, HeLa cells were transfected with siRNAs or siNC and plated on a glass-bottom dish (Mat-Tek). After 24 h, cells were transfected with GFP-SUN2. Experiments were performed on a confocal microscope (FV-1000; Olympus) with a PlanApoN 60 (NA=1.4) oil-immersion objective. After 2 images were obtained (0.2% 488 nm laser transmission), a rectangular region of the nuclear envelope was bleached (100% 488 nm laser transmission), and a further 60 images (10 s intervals) were collected using the original settings. Fluorescence intensity was measured using Metamorph (Molecular Devices, San Jose, CA, USA).

### Plasmids

V5-tagged amphiregulin was previously described [55]. An expression vector for dominant negative KASH (DN-KASH)-mCherry fusion protein was a gift from Daniel Conway (Virginia Commonwealth University, Addgene plasmid #125553) [56]. GFP-tagged mouse Suv39h1 (NM_011514) was purchased from Origene (#MG206488; Rockville, MD, USA). pEGFP-C1-human SUV39H1 was previously described [7]. Point-mutated SUV39H1 (pEGFPC1-Y67A-SUV39H1) was generated by site-directed mutagenesis using appropriate oligo-DNA primer sets and inserted into pEGFP-C1. All cDNA constructs were verified by DNA sequencing.

### Wound healing assay

Details for the procedure were previously described [7]. In brief, as shown in Fig 1, chaetocin (200 nM) was added to the culture medium of confluent MDA-MB-231 cells. The cells were then scratched using a pipette tip. After 5 h incubation, the cells were fixed and labeled for immunofluorescent microscopy. As shown in Fig 5, confluent MDA-MB-231 cells were scratched using a pipette tip (time=0). After incubation for the indicated time, phase-contrast images were obtained using a microscope CKX31 (Olympus, Tokyo, Japan) with a camera (BC-DKM01; bio medical science, Tokyo, Japan). For quantitative analysis, the area of the wound was measured and the ratio to the original wound area is shown. Results are expressed as the mean of three experiments.

### Statistical analysis and quantification of the Golgi dispersion

For Golgi dispersion analysis, experiments were performed at least three times and GM130 staining areas were analyzed using ImageJ (version 1.52). The grey scale images were binarized and the areas of GM130 staining per cell were measured. Results were expressed as means with standard deviations. Statistical significance was assessed using 2-tailed Student’s t-tests.

## Abbreviations

DRB: 5,6-dichloro-1-β-ribofuranosylbenzimidazole
GM130: Golgi matrix protein 130
H3K9me3: histone H3 lysine 9 trimethylation
KASH: Klarsicht, ANC-1, and Syne homology
LINC: linker of nucleoskeleton and cytoskeleton
MT: microtubule
RT-PCR: real-time PCR
siRNA: small interfering RNA
SUN: Sad1 and UNC-84

## Acknowledgements

We thank Yu Nishioka and Junko Imada (Osaka University) for technical assistance and Dr. Tsutomu Kurokawa (Wako Pure Chemical) and Dr. Shuji Matsuura (Osaka University) for valuable discussions.

## Author contributions

Conceptualization: Miki Matsumura, Hiromasa Imaizumi, Nariaki Matsuura

Investigation: Miyu Nishino, Hiromasa Imaizumi, Yuhki Yokoyama, Jun Katahira, Hiroshi Kimura

Supervision: Nariaki Matsuura, Miki Matsumura

Funding acquisition: Nariaki Matsuura, Miki Matsumura

Writing-original draft: Miki Matsumura

Writing-review&editing: Miki Matsumura, Jun Katahira

## Supporting information

**S1 Fig. SUV39H1 associates with the morphology of the Golgi complex. A.** The indicated siRNA pools (three of siRNAs for SUV39H1, four of siRNAs for SETDB1 and SETDB2) were transfected and the cells were stained with DAPI (red) and anti-GM130 mAb (green). Bar, 20 μm. **B.** mRNAs were obtained from SETDB1 or SETDB2 knocked down cells and analyzed by RT-PCR. **C.** Cells were transfected with each of three siRNA against SUV39H1, and the cells were stained with DAPI (red) and anti-GM130 mAb (green). **D.** Cells were transfected with siSUV39H1 or siNC. The cells were then stained with anti-GM130 pAb and anti-Golgi 58K protein, a medial-Golgi marker protein, mAb. Bar, 10 μm.

**S2 Fig. SUV39H1 depletion impairs Golgi complex integrity but does not affect MT or membrane trafficking. A.** Cells were transfected with siSUV39H1. After 36 h incubation, the cells were again transfected with V5-tagged amphiregulin, which is a type I plasma membrane protein. The cells were then fixed and stained with anti-V5 mAb. Bar, 10 μm. **B.** Cells were transfected with siSUV39H1 and stained with anti-β-tubulin mAb (red) and anti-GM130 pAb (green). Bar, 10 μm. **C.** SUV39H1 knock down cells were stained with anti-centrin mAb (red), anti-GM130 pAb (green), and DAPI (blue). Bar, 10 μm. **D.** Cells were incubated with chaetocin (300 nM) in the absence or presence of DRB (50 μM) for 4 h. The cells were then fixed and stained with anti-GM130 mAb and DAPI. Bar, 10 μm.

**S2 Fig. SUN1 protein expression in the SUV39H1-depleted cells.** Whole western blotting shown in Figure 3E indicates SUN1 and β-tubulin protein expression in the SUV39H1 knock down cells.

**S1Table**

**A.** List of siRNAs used in this article. **B.** List of primer sets used in this article.

